# Use of human AML cells to study graft-versus-leukemia immunity in xenogeneic mouse models of GVHD

**DOI:** 10.1101/2024.04.30.591828

**Authors:** Charline Faville, Bianca E Silva, Frédéric Baron, Grégory Ehx

## Abstract

Allogeneic hematopoietic cell transplantation (allo-HCT) is the main therapeutic approach for patients with high-risk acute myeloid leukemia (AML), but the rate of relapse remains high and is associated with poor outcomes. Discovering new approaches to maximize the graft-versus-leukemia (GVL) effects while mitigating graft-versus-host disease (GVHD) should therefore be pursued. Because of the difficulties in modeling AML in mice, patient-derived xenotransplantations (PDX) in immunodeficient NSG mice are preferred to study the GVL effects. In PDX, AML is typically induced through the intravenous injection of cell lines or leukemic blasts obtained from patients. GVHD and GVL effects are induced by (co)-injecting human T cells or peripheral blood mononuclear cells (PBMCs). While this approach enables the induction of systemic leukemia, notably developing in the spleen and bone marrow of the animals, it can also be associated with difficulties in monitoring the disease, notably by flow cytometry. This can be circumvented by using luciferase-expressing AML cells or transplanting the leukemic cells in Matrigel to generate solid tumors that are easier to monitor. Here, we provide detailed instructions on how to prepare human PBMCs and leukemic cells, transplant them, and monitor the disease in NSG mice.

## 1. INTRODUCTION

Acute myeloid leukemia (AML) is an aggressive malignancy characterized by uncontrolled proliferation of abnormal myeloid progenitor cells (blasts) of the hematopoiesis. Currently, AML is the most frequent and lethal leukemia among adults; while chemotherapy results in high remission rates, ∼75% of patients relapse, resulting in an average five-year survival rate of only 40–45% in young patients and less than 10% in the elderly^1^. Following the first cycle of chemotherapy, intermediate- and high-risk AML patients often receive allogeneic hematopoietic cell transplantation (allo-HCT), the only curative option currently available^2, 3^. This therapy relies on two key steps: (1) administration of high doses of chemotherapy to eradicate most leukemic cells and induce medullary aplasia and (2) infusion of a combination of stem cells and mature hematopoietic cells from a genetically different healthy donor. While stem cells colonize the patient’s bone marrow and restore healthy hematopoiesis, mature immune cells recognize and eradicate residual leukemic cells that survived chemotherapy, a process known as the graft-versus-leukemia (GVL) effect. Relapse prevention by the GVL effect is mostly mediated by transplanted immune cells that recognize leukemic cells as “foreign” due to the HLA mismatch between donor and recipient and/or the presentation of MHC peptides derived from polymorphic genomic regions that differ between donor and recipient^4^. However, this allorecognition can also cause the donor’s immune cells to attack the recipient’s healthy tissues, resulting in an inflammatory condition known as Graft-versus-Host Disease (GVHD) that can lead to death^5^. Therapies aimed at improving the GVL effects while mitigating GVHD are therefore under intense investigation^6^.

The currently accepted AML pathogenesis mechanism describes that the malignant transformation of hematopoietic progenitors results from the effect of one or several cooperating driver mutations^7, 8^. Therefore, modeling AML in rodents relies on engineering these mutations in the germline or retroviral transduction of bone marrow cells followed by transplantation. Apart from the obvious species-specific difference, these models are limited because engineered genetic lesions only partially mimic those found in the leukemic blasts of patients, as – even in the presence of leading cytogenetic lesions – AML blasts often contain one or several potentially cooperating mutations^9^. Therefore, using patient-derived xenotransplantations (PDX) or cell line-derived xenotransplantations (CDX) in immunodeficient NOD-scid-IL2rg^null^ (NSG) mice is preferred when studying AML *in vivo*^10^.

Over the last decade, our lab has extensively studied the pathogenesis and effects of multiple treatments on xenogeneic GVHD, either alone^11-15^ or on both GVHD and GVL effects^16-18^. Our experience with these models highlighted two main technical limitations needing special attention when performing such experiments.

The first is the capacity of primary AML cells to engraft. Indeed, this can be limited by the presence of mature T- or B cells in the transplanted AML sample that mediate GVHD, thereby limiting AML engraftment and/or leading to the death of the mice before AML angreftment^19^. Therefore, T- and B cells should be depleted from primary AML cells, or cell lines should be used. Alternatively, in experiments only focused on GVL effects and when T- and B cells cannot be depleted, NSG mice lacking mouse MHC molecules can be used as they develop GVL effects without GVHD^20^. Here, it is worth mentioning that an extended period is sometimes needed to reach detectable engraftment of leukemic cells when using some primary cells or cell lines. This has caused several investigators to conclude that some AML samples cannot engraft in NSG mice (six weeks post-transplantation), while others demonstrated that by using an extended follow-up time (several months), virtually all AML cells eventually engraft^21^. Therefore, patience, in addition to T-/B cell depletion, can be the key to engrafting AML cells. Finally, NSG mice expressing SCF, GM-CSF, and IL-3 (NSG-S) can be used to boost the engraftment of AML cells^22^.

The second is the capacity to detect AML cells to monitor either the engraftment or the GVL effects. We observed that T cells are the main immune cell population engrafting in NSG mice upon intravenous infusion of peripheral blood mononuclear cells (PBMCs)^12^. Consequently, the detection of AML cells could be based on flow cytometry analyses of the CD33^+^ population in the blood of mice transplanted with AML cells only or in combination with PBMCs. However, such analyses are limited by the capacity of AML cells to exit the bone marrow and be mobilized to the peripheral blood^23, 24^ and by the expression of a specific marker, such as CD33, by leukemic cells. Indeed, only 75% of patients express this marker and the subset of cells that regenerate leukemia in immunocompromised mice does not tend to express CD33^25^. Consequently, using luciferase-expressing cell lines or Matrigel-based subcutaneous injections of primary cells (generating measurable tumors) may be needed to enable the monitoring of GVL effects^12^.

In the present work, we provide detailed instructions on how to prepare human PBMCs and leukemic cells, transplant them, and monitor the disease in NSG mice.

## 2. MATERIALS

Ensure that all reagents intended for injection into mice remain sterile. Prepare and store all reagents at room temperature (unless indicated otherwise).

### 2.1. Cell line preparation for transplantation in NSG mice

1. AML cell line, expressing luciferase (OCI-AML3-luc, Ubigene, YC-C022-Luc-P) or not (OCI-AML3, DSMZ, Ref. ACC 582).
2. Phosphate-buffered saline, pH 7.4 (PBS, Gibco, Ref. 14190-094).
3. Trypan blue (Sigma, Ref. T8154).
4. Dual-chamber counting slides for cell counter (Biorad, 145-0011).
5. Automatic cell counter (Biorad, TC-20, Ref. 1450102).
6. Sterile FACS tube (Falcon, Ref. 352063).
7. Cell strainer, 70 µm (VWR, Ref. 732-2758).

### 2.2. Preparation of primary blasts for transplantation in NSG mice

1. Red blood cell lysis buffer (5X): 8.28 g of ammonium chloride (Merck, Ref. 1.01145.0500), 0.036 g of EDTA (Merck, Ref. 1.08418.1000), and 1 g of potassium bicarbonate (Merck, Ref. 1.04854.0500) in 1 L of deionized water. Adjust to pH 7.1-7.2 and sterilize by autoclaving (*see* **Note 1**).
2. Supplemented RPMI (Gibco, Ref. 21875034): 10% heat-inactivated fetal bovine serum (FBS, Gibco, Ref. A5256701), 1% penicillin-streptomycin (Gibco, Ref. 15140-122) in RPMI, pre-heat at 37°C.
3. Sterile FACS tubes (Falcon, Ref. 352063).
4. Falcon 50 mL conical tubes (Falcon, Ref. 352098).
5. Hematological cell counter (Sysmex, XN-450).
6. PBS (Gibco, Ref. 14190-094).
7. Flow cytometry staining buffer: 3% FBS in PBS.
8. Depletion assessment mix of antibodies for flow cytometry staining: 2 µL of anti-CD4-FITC (BD Biosiences, Ref. 345768), 1 µL of anti-CD33-PE (Invitrogen eBioscience, Ref. 12-0339-42), 1 µL of anti-CD19-APC (Invitrogen eBioscience, Ref. 17-0199-42), 1 µL of anti-CD3-V450 (BD Biosiences, Ref. 560365), 1 µL of anti-CD8-APC-Cy7 (BD Biosiences, Ref. 557834), 0.5 µL of anti-CD45-BV510 (BD Biosiences, Ref. 563204), 1 µL of anti-CD117-APC-R700 (BD Biosiences, Ref. 565195), 1 µL of anti-CD56-Pe-Cy7 (Sony, Ref. 2191590) and 1 µL of anti-CD34-PerCP-Cy5.5 (BD Biosiences, Ref. 347222) per tube to label.
9. Human Fc Blocker (Biolegend, Ref. 422302).
10. EasySep Human Biotin Positive Selection Kit II (Stemcell Technologies, Ref. 17663).
11. Anti-CD3-biotin (Biolegend, Ref. 344820).
12. Anti-CD19-biotin (Biolegend, Ref. 302204).
13. Non-graduated polystyrene culture tubes - 14 mL - 17 mm (Roth®, ref. AEX9.1).
14. Cell strainer, 70 µm (VWR, Ref. 732-2758).
15. Flow cytometer equipped with violet (405 nm) and red (561 nm) lasers, and 780/60 and 450/40 nm bandpass filters (Becton Dickinson, LSR Fortessa).
16. Water bath at 37°C.
17. EasySep magnet (Stemcell Technologies, Ref. 18001).
18. Anti-CD3-V450 (Becton Dickinson, Ref. 560365).
19. Anti-CD19-APC-Cy7 (Becton Dickinson, Ref. 55779).
20. Hematological cell counter (Sysmex, XN-450).

### 2.3. PBMCs isolation from healthy volunteers

1. PBS (Gibco, Ref. 14190-094).
2. Ficoll-Paque PLUS (GE Healthcare, Ref. 17144003).
3. Falcon 50 mL conical tubes (Falcon, Ref. 352098).
4. Hematological cell counter (Sysmex, XN-450).
5. Cell strainer, 70 µm (VWR, Ref. 732-2758).
6. Sterile FACS tubes (Falcon, Ref. 352063).

### 2.4. Transplantation to NSG mice

1. NSG mice (Jackson Laboratory, Ref. 005557).
2. Antibiotics: 0.1% Baytril 25 mg/mL (Bayer, Ref. 0369124) in autoclaved water.
3. Matrigel matrix (Corning, Ref. 354234) (*see* **Note 2**).
4. BD Plastipak Syringe (BD Biosiences, Ref. 303172).
5. BD Microlance 3-26G x ½” needle for syringe (BD Biosiences, Ref. 560365).
6. PBS (Gibco, Ref. 14190-094).
7. BD Plastipak insulin syringe with Micro-Fine + crimped needle (BD Biosiences, Ref. 324891)

### 2.5. Monitoring AML engraftment by flow cytometry

1. PBS (Gibco, Ref. 14190-094).
2. Flow cytometry staining buffer: 3% FBS in PBS.
3. Mix of antibodies for flow cytometry staining: 1 µL of anti-CD33-PE (Invitrogen eBioscience, Ref. 12-0339-42) and 0.5 µL of anti-CD45-BV510 (BD Biosiences, Ref. 563204) per tube.
4. Human Fc Blocker (Biolegend, Ref. 422302).
5. Red blood cell (RBC) lysis buffer 10X (Invitrogen eBioscience, Ref. 00430054): dilute 10- fold with deionized water.
6. Heparized capillaries (VWR, Ref. 521-9100).
7. EDTA Microtainer tubes (Becton Dickison, Ref. 365975).
8. Flow cytometry polystyrene tubes (Corning, Ref. 352008).
9. Mouse restraining device (Midsci, Ref. MH-100).
10. BD Microlance 3-26G x ½” needle for syringe (BD Biosiences, Ref. 560365).
11. Flow cytometer with violet (405 nm) and blue (488 nm) lasers and 585/42 and 525/50 nm bandpass filters (Becton Dickinson, LSR Fortessa).
12. Hematological cell counter (Sysmex, XN-450).
13. Antiseptic solution: 70 % ethanol (Merck, Ref. 100986) in desionized water.

### 2.6. Monitoring the engraftment by bioluminescence imaging

1. Beetle luciferin, potassium salt (Promega, Ref. E1605).
2. PBS (Gibco, Ref. 14190-094).
3. Isoflurane 100% (Isoflo, Zoetis, Ref. 5414736033471).
4. BD Plastipak insulin syringe with Micro-Fine + crimped needle (BD Biosiences, Ref. 324891).
5. Polypropylene microcentrifuge tubes, 1.5 mL (Greiner, Ref. 616201).
6. IVIS Spectrum Imaging system (Revvity, Ref. 124262).

### 2.7. Monitoring subcutaneous tumor size with calipers

1. Calipers (VWR, Ref. 62379-531).

## 3. METHODS

All cell preparations are carried out in Class II Biological Safety Cabinets (BSL2) to maintain the sterility of the samples before transplanting mice and to protect the operator from possible pathogens present in human blood samples. Diligently follow all waste disposal regulations when disposing of biological waste and materials. Wear protective gloves and a lab coat at all steps of the following protocols.

### 3.1. Cell line preparation for transplantation into NSG mice

1. Combine 20 µL of the AML cell line suspension (either luciferase expressing or not) with 20 µL of Trypan Blue and mix well by gently pipetting up and down ten times. Within 5 min of mixing, pipet 10 µL of the mixture into the outer opening of either of the two chambers of the counting slide. Insert the slide into the slot of the cell counter to activate cell counting.
2. Based on the cell count obtained, collect the appropriate volume to inject 1 million cells per mouse to be transplanted. Consider preparing additional cells beyond the exact requirement to accommodate potential sample loss during the injection process. Typically, allocate one dead volume for every three mice to be transplanted.
3. Centrifuge the cell suspension at 500 x *g* for 5 min at room temperature and carefully remove the supernatant with a pipette.
4. Resuspend the cells at a concentration of 10 million cells per mL with PBS.
5. Filter the suspension with a 70 µm cell strainer into a sterile FACS tube.
6. Keep tubes on ice until transplantation.

### 3.2. Preparation of primary blasts for transplantation into NSG mice

1. Place 10 mL of peripheral blood or bone marrow aspirate into a 50 mL Falcon tube and add four volumes of 5X sterile red blood cell lysis buffer (*see* **Note 3**). Homogenize the tube by inversion immediately after adding the lysis buffer.
2. Place the tube on a rocker set to 25 oscillations per minute and incubate for 15 min.
3. Centrifuge the tube at 300 x *g* for 10 min at room temperature. Carefully remove the supernatant with a pipette, leaving 1-2 mL of supernatant above the pellet, and resuspend the pellet with a micropipette.
4. Add 5 mL of supplemented RPMI (*see* **Note 4**).
5. Centrifuge the tube at 500 x *g* for 5 min at room temperature. Carefully remove as much supernatant as possible with a pipette. Resuspend the pellet in 5 mL of supplemented RPMI.
6. Count the cells with the hematological cell counter and record the white blood cell (WBC) counts.
7. Keep 1 million cells in a FACS tube to evaluate the depletion efficacy of B- and T cells (*see* step 23).
8. Adjust the remaining cells to 100 million WBCs/mL with flow cytometry staining buffer, filter the cell suspension with a 70 µm cell strainer, and transfer the cells to a 14 mL non-graduated polystyrene culture tube.
9. Add the anti-CD3-biotin and anti-CD19-biotin antibodies at a concentration of 0.3 - 3 µg/mL of sample. Incubate for 20 min at room temperature.
10. Top up to 10-fold excess FACS staining. Centrifuge at 300 x *g* for 10 min at room temperature with a low brake. Carefully remove the supernatant with a pipette and resuspend the pellet in the same volume of PBS as in step 8.
11. Add the “Selection Cocktail” (from the EasySep Human Biotin Positive Selection Kit II) to the sample at a concentration of 100 µL per mL of sample (do not vortex the cocktail before adding it to the tube). Mix and incubate for 15 min at room temperature.
12. Vortex the tube of “RapidSpheres” (from the EasySep Human Biotin Positive Selection Kit II) for 30 seconds.
13. Add the “RapidSpheres” to the sample at a concentration of 50 µL per mL of sample. Mix and incubate for 10 min at room temperature.
14. Top up the tube to 10 mL with PBS. Homogenize gently with a pipette.
15. Place the tube (without the cap) into the EasySep magnet. Incubate for 5 min at room temperature.
16. After the incubation, keep the tube inside the magnet and carefully tilt the magnet to pour off the supernatant into a new tube. Discard the first tube and place a new tube into the magnet.
17. Repeat steps 15 and 16 four times to perform a total of five incubations in the magnet to ensure complete depletion. After the last incubation, pour the supernatant into a new tube.
18. Count the WBCs with the hematological cell counter.
19. Centrifuge the cell suspension at 500 x *g* for 5 min at room temperature. Carefully remove the supernatant with a pipette.
20. Resuspend the cells at a concentration of 10 million cells per mL in PBS.
21. Filter the suspension with a 70 µm sterile cell strainer into a new tube.
22. Transfer 1 million cells into a FACS tube and keep the remaining cells on ice until transplantation.
23. Centrifuge the 1 million cells from steps 7 and 22 for 5 min at 500 x *g*.
24. Discard the supernatant, resuspend the cells in 1 mL of flow cytometry staining buffer, and centrifuge for 5 min at 500 x *g* at room temperature.
25. Discard the supernatant and resuspend the cells in 50 µL of staining buffer.
26. Add 5 µL of Fc Blocker to each tube, vortex, and incubate for 10 min at room temperature.
27. Add 9.5 µL of depletion assessment antibody mix to each tube, vortex, and incubate for 20 min at 4°C (*see* **Note 5**).
28. Add 1 mL of PBS to each tube, vortex, and centrifuge for 5 min at 500 x *g*.
29. Resuspend the cells in 200 µL of PBS and analyze the B/T-cell depletion efficacy as well as the blast frequency in the sample to be transplanted into the mice (Figure 1A-B).

**Figure 1.**
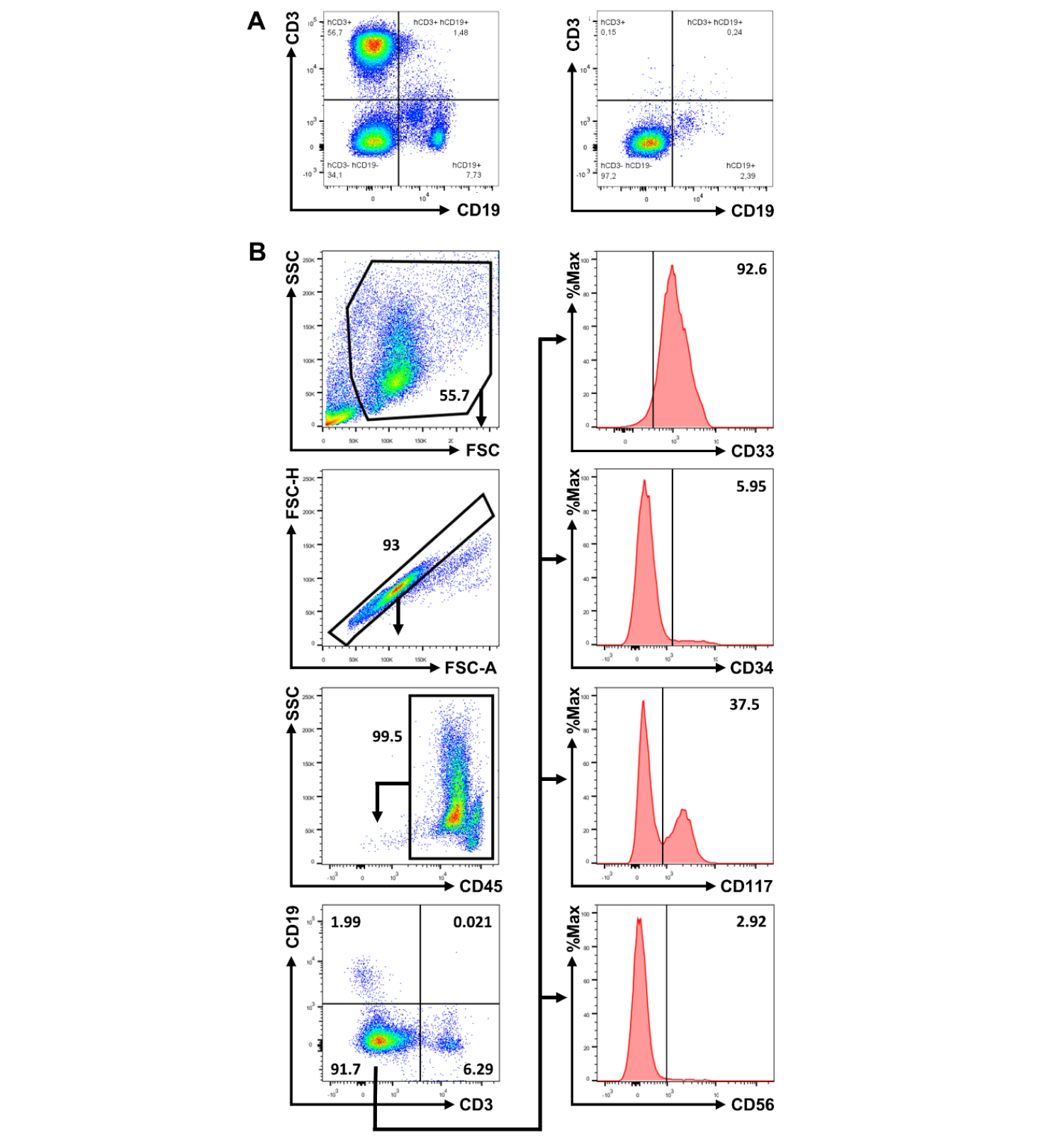
**(A)** Depletion of B- and T cells from primary AML samples with the EasySep Human Biotin Positive Selection Kit II. The plot on the left corresponds to the cells from step 7 (before depletion), while the plot on the right corresponds to the cells from step 22 (after the depletion). **(B)** Gating strategy for the analysis of the phenotype of AML blasts in primary samples by flow cytometry.

### 3.3. PBMCs isolation from healthy volunteers

Here, it is assumed that PBMCs are injected simultaneously with AML cells. Alternatively, PBMCs can be transplanted upon observing that AML cells have reached the desired level of engraftment. This is especially relevant when using primary samples, which can take several weeks to reach detectable engraftment levels. Delaying the PBMC transplantation can prevent the development of lethal GVHD (occurring typically within 3-6 weeks post-transplantation^12^) before the engraftment of AML cells. Finally, lower numbers of PBMCs can be transplanted by sub-lethally irradiating the mice 24h before transplantation (*see* step 9).

1. If working with buffy coats from healthy volunteers, transfer 17.5 mL of cell suspension into a 50 mL Falcon tube and add 17.5 mL of PBS, mix by inversion. Otherwise, proceed to step 2.
2. Transfer 15 mL of Ficoll-Paque into a 50 mL Falcon tube.
3. Carefully overlay 35 mL of peripheral blood or diluted buffy coat onto the Ficoll-Paque. Important: When overlaying the sample, do not mix the Ficoll-Paque with the diluted blood sample.
4. Centrifuge for 20 min at 500 x g at room temperature with no brakes.
5. With a pipet, discard ∼10 mL of plasma from the top layer and collect the mononuclear cells (PBMCs), located at the interface between the Ficoll and the plasma (thin white layer right under the plasma), into a new 50 mL Falcon tube.
6. Adjust the volume of PBMCs to 50 mL with PBS and centrifuge for 5 min at 500 x *g* at room temperature.
7. Discard the supernatant and resuspend the cells in 5 mL of PBS.
8. Count the cells with the hematological cell counter.
9. Adjust the WBC concentration to 200 million cells per mL. If mice have been irradiated with 2 Gy 24h before the transplantation, PBMCs should be adjusted to 20 million per mL.
10. Filter the suspension with a 70 µm sterile cell strainer into a new sterile tube.
11. Keep the cells on ice until transplantation.

### 3.4. Transplantation into NSG mice

NSG mice are typically transplanted between the ages of 8 to 11 weeks. As NSG mice are highly immunodeficient, supplement their drinking water with antibiotics (Baytril) to reduce their risk of developing infections during the experiment.

1. For intravenous transplantation, either dilute the AML cells 2-fold with PBS to reach a final concentration of 5 million cells per mL or combine the AML cells with PBMCs at a 1:1 ratio to reach a final concentration of 5 million AML cells per mL and 100 million PBMCs per mL.
2. For subcutaneous injections, dilute the AML cells 2-fold with Matrigel and mix well by pipetting up and down. Keep the cells on ice until injection.
3. For intravenous transplantations, inject 200 µL of suspension into the tail vein of each mouse (1 million AML cells + 20 million PBMCs/mouse, or 1 million AML cells + 2 million PBMCs/mouse if mice have been irradiated) with a BD Microlance 3 syringe.
4. For subcutaneous transplantations, inject 200 µL of the AML cells + Matrigel suspension (1 million AML cells per animal) into the flank of each mouse with an insulin syringe.
5. If subcutaneous AML transplantations are being performed, dilute the PBMCs 2-fold with PBS and inject 200 µL of suspension into the tail vein of each mouse (20 million PBMCs per mouse or 2 million if animals have been irradiated).
6. Once the AML cells and/or PBMCs have been transplanted, the animals might develop symptoms of AML and/or GVHD. These symptoms include abnormal weight change (increasing or decreasing), anemia, hunching, and reduced activity. Score the animals every 2-3 days for these symptoms (two grades of severity for each symptom, with a weight change of 10% = grade 1 and a weight change of 20% = grade 2). Sacrifice all animals reaching a score of 6/8, having a tumor size of 1 cm³, or experiencing a weight change greater than 20%. Thoroughly follow the guidelines of your local ethics committee.

### 3.5. Monitoring the AML engraftment by flow cytometry

Starting from seven days post-transplantation, the engraftment level of AML cells can be monitored in the peripheral blood of NSG mice to evaluate the GVL effects. A low engraftment (frequency of AML cells in peripheral blood) could reflect greater GVL effects. This procedure can also be used to monitor the evolution of the AML engraftment in mice that did not receive PBMCs simultaneously with AML cells.

1. Restrain the mouse using the restraint device with the tail protruding.
2. Disinfect the tail with an antiseptic solution.
3. Heat the tail with an infrared lamp for 2-3 min to increase blood flow in the tail and induce dilatation of the veins.
4. Puncture the tail vein with the needle of a syringe and collect the blood with a capillary (one capillary per mouse is sufficient).
5. Transfer the blood to a Microtainer tube and return the mouse to its cage.
6. Count the WBCs with the hematological counter.
7. Transfer the remaining blood (max 50 µL) into a FACS tube and add 1 mL of 1X red blood cell lysis buffer, vortex well, and incubate for 5 min at room temperature.
8. Add 1 mL of PBS to each tube, vortex, and centrifuge for 5 min at 500 x *g* at room temperature.
9. Discard the supernatant, add 1 mL of staining buffer, vortex, and centrifuge for 5 min at 500 x *g* at room temperature.
10. Discard the supernatant, resuspend the cells in 50 µL of staining buffer, and add 5 µL of Fc Blocker to each tube. Vortex and incubate for 10 min at room temperature.
11. Add 1 µL of each antibody (anti-CD45 and anti-CD33). Vortex and incubate for 20 min at 4°C.
12. Add 1 mL of PBS to each tube, vortex, and centrifuge for 5 min at 500 x *g* at room temperature.
13. Discard the supernatant and resuspend the cells in 200 µL of PBS.
14. Keep the cells at 4°C until analysis by flow cytometry (Figure 2).
15. Based on the frequency of CD33^+^ cells among CD45^+^ cells (WBCs), and the concentration of WBCs obtained with the hematological counter, compute the concentration of CD33^+^ cells in the blood of the animals. Since normal granulocytes and monocytes do not engraft in NSG mice, the frequency of CD33^+^ cells can be used as a proxy of the engraftment of AML cells.

**Figure 2.**
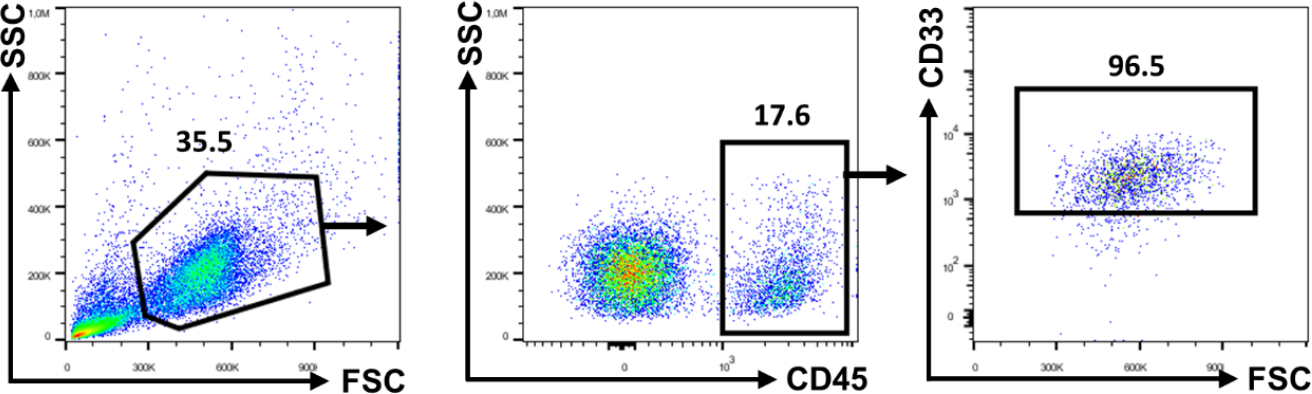
Gating strategy for the analysis of AML cells (defined as human CD45^+^CD33^+^ cells) in the blood of NSG mice transplanted intravenously.

### 3.6. Monitoring the engraftment by bioluminescence imaging

If luciferase-expressing AML cells have been transplanted into mice, *in vivo* bioluminescence imaging can be used to monitor the engraftment of AML cells. The advantage of this technique is that all AML cells (not only those in circulation and/or expressing CD33) can be detected.

1. Reconstitute luciferin in PBS to a concentration of 15 mg/mL. Homogenize well. The luciferin solution can be aliquoted and kept at -80°C.
2. Initialize the Xenogen IVI Spectrum following the specific instructions of the device.
3. Saturate the inhalation chamber with isoflurane.
4. Subcutaneously inject 200 µL of luciferin solution into a maximum of 5 mice at a time.
5. Return the mice to their cages and wait for 8 min.
6. Place the mice into the inhalation chamber for 3 min. Then, place the mice into the IVIS with isoflurane masks, keeping them under anesthesia.
7. Capture an initial image with the acquisition time set at 1 minute, then adjust the acquisition time accordingly: Decrease if the signal is oversaturated or increase if the signal is absent.
8. Image analysis can be performed with Living Image software. The light intensity for each tumor or across the whole body of the animal (if intravenous transplantation was performed) is proportional to the engraftment of AML cells (Figure 3).

**Figure 3.**
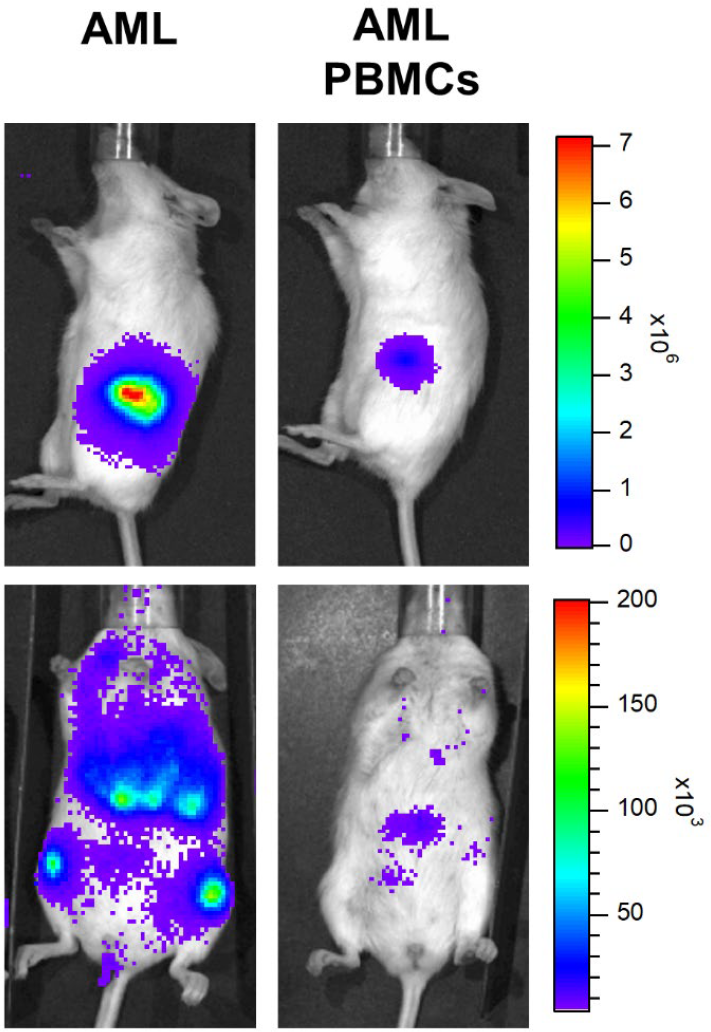
NSG mice were transplanted with AML cells (THP-1-luc) either subcutaneously in matrigel (top) or intravenously (bottom). Mice received PBMCs (right) or not (left). Bioluminescence imaging (photons/sec) shows that the presence of PBMCs reduces the growth of AML cells (GVL effects).

### 3.7. Monitoring subcutaneous tumor size with calipers

If AML cells (either expressing luciferase or not) are transplanted subcutaneously, tumor size can be monitored with calipers.

1. Securely hold the tail of the mouse betwen your outermost and middle fingers, grasp the loose skin at the nape of the neck between your forefinger and thumb, and align the animal in your hand to expose the tumor.
2. With a caliper, measure the length (L) and width (W) of the tumor.
3. Assuming that the tumor is an ellipsoid, use the following formula to compute its volume: V_T_ = 0.5 × L × W².

## 4. Notes

1. Store the stock lysis buffer at 4°C and protect it from light. Pre-warm the buffer to room temperature before using it to lyse red blood cells.
2. Matrigel is preserved at -20°C but has to be thawed at 4°C 24h before transplantation. Keep thawed Matrigel on ice.
3. If necessary, and if enough blood or bone marrow aspirate is available, prepare as many 50 mL tubes as needed. A peripheral blood volume of 50 mL from an AML patient with active disease is typically sufficient to transplant 10 mice.
4. If a signficant number of red blood cells remain visible after the initial lysis, a second lysis can be performed.
5. Alternatively, the cells can be incubated overnight at 4°C to maximize the staining efficacy^26^.

## Acknowledgment

This work was supported by the FNRS Belgium, the Leon Fredericq Foundation, and the ME-TO-YOU Foundation. CF and BES are research fellows at the FNRS. G.E. and FB are research associates at the FNRS.

## References

1. Craddock, C., Tauro, S., Moss, P. & Grimwade, D. Biology and management of relapsed acute myeloid leukaemia. 2005 Br J Haematol 129, 18–34.

2. Baron, F. & Storb, R. Allogeneic hematopoietic cell transplantation as treatment for hematological malignancies: a review. 2004 Springer Seminars in Immunopathology 26, 71–94.

3. Döhner, H., Wei, A.H., Appelbaum, F.R., Craddock, C., DiNardo, C.D., Dombret, H., Ebert, B.L., Fenaux, P., Godley, L.A., Hasserjian, R.P., Larson, R.A., Levine, R.L., Miyazaki, Y., Niederwieser, D., Ossenkoppele, G., Röllig, C., Sierra, J., Stein, E.M., Tallman, M.S., Tien, H.F., Wang, J., Wierzbowska, A. & Löwenberg, B. Diagnosis and management of AML in adults: 2022 recommendations from an international expert panel on behalf of the ELN. 2022 Blood 140, 1345–1377.

4. Fontaine, P., Roy-Proulx, G., Knafo, L., Baron, C., Roy, D.C. & Perreault, C. Adoptive transfer of minor histocompatibility antigen-specific T lymphocytes eradicates leukemia cells without causing graft-versus-host disease. 2001 Nat Med 7, 789–794.

5. Zeiser, R., Socie, G. & Blazar, B.R. Pathogenesis of acute graft-versus-host disease: from intestinal microbiota alterations to donor T cell activation. 2016 Br J Haematol 175, 191–207.

6. Servais, S., Beguin, Y., Delens, L., Ehx, G., Fransolet, G., Hannon, M., Willems, E., Humblet-Baron, S., Belle, L. & Baron, F. Novel approaches for preventing acute graft-versus-host disease after allogeneic hematopoietic stem cell transplantation. 2016 Expert Opin Investig Drugs 25, 957–972.

7. Grossmann, V., Schnittger, S., Kohlmann, A., Eder, C., Roller, A., Dicker, F., Schmid, C., Wendtner, C.-M., Staib, P. & Serve, H. A novel hierarchical prognostic model of AML solely based on molecular mutations. 2012 Blood, The Journal of the American Society of Hematology 120, 2963–2972.

8. Rose, D., Haferlach, T., Schnittger, S., Perglerová, K., Kern, W. & Haferlach, C. Subtype-specific patterns of molecular mutations in acute myeloid leukemia. 2017 Leukemia 31, 11–17.

9. Papaemmanuil, E., Döhner, H. & Campbell, P.J. Genomic Classification in Acute Myeloid Leukemia. 2016 N Engl J Med 375, 900–901.

10. Fisher, J.N., Kalleda, N., Stavropoulou, V. & Schwaller, J. The Impact of the Cellular Origin in Acute Myeloid Leukemia: Learning From Mouse Models. 2019 Hemasphere 3, e152.

11. Courtois, J., Ritacco, C., Dubois, S., Canti, L., Vandenhove, B., Seidel, L., Daulne, C., Caers, J., Servais, S., Beguin, Y., Ehx, G. & Baron, F. Itacitinib prevents xenogeneic GVHD in humanized mice. 2021 Bone Marrow Transplant 56, 2672–2681.

12. Ehx, G., Somja, J., Warnatz, H.J., Ritacco, C., Hannon, M., Delens, L., Fransolet, G., Delvenne, P., Muller, J., Beguin, Y., Lehrach, H., Belle, L., Humblet-Baron, S. & Baron, F. Xenogeneic Graft-Versus-Host Disease in Humanized NSG and NSG-HLA-A2/HHD Mice. 2018 Front Immunol 9, 1943.

13. Hannon, M., Lechanteur, C., Lucas, S., Somja, J., Seidel, L., Belle, L., Bruck, F., Baudoux, E., Giet, O., Chantillon, A.M., Delvenne, P., Drion, P., Beguin, Y., Humblet-Baron, S. & Baron, F. Infusion of clinical-grade enriched regulatory T cells delays experimental xenogeneic graft-versus-host disease. 2014 Transfusion 54, 353–363.

14. Grégoire, C., Ritacco, C., Hannon, M., Seidel, L., Delens, L., Belle, L., Dubois, S., Vériter, S., Lechanteur, C., Briquet, A., Servais, S., Ehx, G., Beguin, Y. & Baron, F. Comparison of Mesenchymal Stromal Cells From Different Origins for the Treatment of Graft-vs.-Host-Disease in a Humanized Mouse Model. 2019 Front Immunol 10, 619.

15. Delens, L., Ehx, G., Somja, J., Vrancken, L., Belle, L., Seidel, L., Grégoire, C., Fransolet, G., Ritacco, C., Hannon, M., Dubois, S., Beguin, Y., Baron, F. & Servais, S. In Vitro Th17-Polarized Human CD4(+) T Cells Exacerbate Xenogeneic Graft-versus-Host Disease. 2019 Biol Blood Marrow Transplant 25, 204–215.

16. Ehx, G., Ritacco, C., Hannon, M., Dubois, S., Delens, L., Willems, E., Servais, S., Drion, P., Beguin, Y. & Baron, F. Comprehensive analysis of the immunomodulatory effects of rapamycin on human T cells in graft-versus-host disease prophylaxis. 2021 Am J Transplant 21, 2662–2674.

17. Ehx, G., Fransolet, G., de Leval, L., D’Hondt, S., Lucas, S., Hannon, M., Delens, L., Dubois, S., Drion, P., Beguin, Y., Humblet-Baron, S. & Baron, F. Azacytidine prevents experimental xenogeneic graft-versus-host disease without abrogating graft-versus-leukemia effects. 2017 Oncoimmunology 6, e1314425.

18. Ritacco, C., Köse, M.C., Courtois, J., Canti, L., Beguin, C., Dubois, S., Vandenhove, B., Servais, S., Caers, J., Beguin, Y., Ehx, G. & Baron, F. Post-transplant cyclophosphamide prevents xenogeneic graft-versus-host disease while depleting proliferating regulatory T cells. 2023 iScience 26, 106085.

19. von Bonin, M., Wermke, M., Cosgun, K.N., Thiede, C., Bornhauser, M., Wagemaker, G. & Waskow, C. In vivo expansion of co-transplanted T cells impacts on tumor re-initiating activity of human acute myeloid leukemia in NSG mice. 2013 PLoS One 8, e60680.

20. Brehm, M.A., Kenney, L.L., Wiles, M.V., Low, B.E., Tisch, R.M., Burzenski, L., Mueller, C., Greiner, D.L. & Shultz, L.D. Lack of acute xenogeneic graft-versus-host disease, but retention of T-cell function following engraftment of human peripheral blood mononuclear cells in NSG mice deficient in MHC class I and II expression. 2019 Faseb j 33, 3137–3151.

21. Paczulla, A.M., Dirnhofer, S., Konantz, M., Medinger, M., Salih, H.R., Rothfelder, K., Tsakiris, D.A., Passweg, J.R., Lundberg, P. & Lengerke, C. Long-term observation reveals high-frequency engraftment of human acute myeloid leukemia in immunodeficient mice. 2017 Haematologica 102, 854–864.

22. Wunderlich, M., Chou, F.S., Link, K.A., Mizukawa, B., Perry, R.L., Carroll, M. & Mulloy, J.C. AML xenograft efficiency is significantly improved in NOD/SCID-IL2RG mice constitutively expressing human SCF, GM-CSF and IL-3. 2010 Leukemia 24, 1785–1788.

23. Lopez-Millan, B., Diaz de la Guardia, R., Roca-Ho, H., Anguita, E., Islam, A., Romero-Moya, D., Prieto, C., Gutierrez-Agüera, F., Bejarano-Garcia, J.A., Perez-Simon, J.A., Costales, P., Rovira, M., Marín, P., Menendez, S., Iglesias, M., Fuster, J.L., Urbano-Ispizua, A., Anjos-Afonso, F., Bueno, C. & Menendez, P. IMiDs mobilize acute myeloid leukemia blasts to peripheral blood through downregulation of CXCR4 but fail to potentiate AraC/Idarubicin activity in preclinical models of non del5q/5q-AML. 2018 Oncoimmunology 7, e1477460.

24. Díaz de la Guardia, R., Velasco-Hernandez, T., Gutiérrez-Agüera, F., Roca-Ho, H., Molina, O., Nombela-Arrieta, C., Bataller, A., Fuster, J.L., Anguita, E., Vives, S., Zamora, L., Nomdedeu, J., Gómez-Casares, M.T., Ramírez-Orellana, M., Lapillonne, H., Ramos-Mejia, V., Rodríguez-Manzaneque, J.C., Bueno, C., Lopez-Millan, B. & Menéndez, P. Engraftment characterization of risk-stratified AML in NSGS mice. 2021 Blood Adv 5, 4842–4854.

25. Vercauteren, S., Zapf, R. & Sutherland, H. Primitive AML progenitors from most CD34(+) patients lack CD33 expression but progenitors from many CD34(-) AML patients express CD33. 2007 Cytotherapy 9, 194–204.

26. Whyte, C.E., Tumes, D.J., Liston, A. & Burton, O.T. Correction: Do more with Less: Improving High Parameter Cytometry Through Overnight Staining. 2023 Curr Protoc 3, e678.

